# Protective function of *ex vivo* expanded CD8 T cells in a mouse model of adoptive therapy for cytomegalovirus infection depends on integrin beta 1 but not CXCR3, CTLA4, or PD-1 expression

**DOI:** 10.1101/2024.03.16.585350

**Authors:** Xiaokun Liu, Rodrigo Gutierrez Jauregui, Yvonne Lueder, Stephan Halle, Laura Ospina-Quintero, Christiane Ritter, Anja Schimrock, Stefanie Willenzon, Anika Janssen, Karen Wagner, Martin Messerle, Berislav Bošnjak, Reinhold Förster

## Abstract

The adoptive transfer of virus-specific T cells (VSTs) represents a therapeutic option for viral infection treatment in immunocompromised patients. Before administration, *ex vivo* culture enables VST expansion. However, it is unclear how *ex vivo* expansion affects the circulation, homing, and intra-tissue migration of administered VSTs. We established a model of VST immunotherapy of acute cytomegalovirus infection using adoptive transfer of *ex vivo* expanded OT-I CD8 T cells (recognizing SIINFEKL peptide) into *Rag2*^-/-^ mice infected with murine cytomegalovirus (MCMV) encoding for the SIINFEKL peptide. *Ex vivo* expansion induced an effector T cell phenotype and affected the expression of integrins and chemokine receptors. CRISPR/Cas9-mediated gene deletions enabled us to address the role of selected genes in the homing of VSTs following intravenous administration. We found that deletion of *Itgb1*, encoding for integrin beta 1, prevented OT-I cells from entering infected organs and drastically reduced their number in blood, suggesting that adoptively transferred VSTs primarily expand in the infected tissues. In contrast, *Cxcr3*^-/-^ OT-I cells provided equal protection as their *Cxcr3*^+/+^ counterparts, indicating that this chemokine receptor does not contribute to VST entry into infected organs. Further, *Pdcd1* and *Ctla4* deletion did not impair the transferred OT-I cells’ ability to protect mice from MCMV, arguing against quick exhaustion of VSTs with an effector T cell phenotype. Together, these data indicate that *ex vivo* expansion affects migration and activation properties of VSTs and suggest that future clinical evaluation of adoptive T cell therapy efficacy should include homing molecule expression assessment.

## 1 Introduction

CD8 T cells are crucial for the defense against intracellular pathogens and immune surveillance against cancer. ^1,2^ Equipped with T cell receptors (TCR), CD8 T cells can specifically and selectively destroy tumor- or pathogen-infected cells by recognizing tumor- or pathogen-specific peptides presented by major histocompatibility complex (MHC) class I molecules on the target cell surface. Given the vast number of potential antigens to which specific responses have to be mounted, it is not surprising that the frequency of T cells expressing TCR with the same antigen specificity is only around one per hundred thousand to a million T cells. ^3^ In a search for antigens, the naïve CD8 T cells generated in thymus traffic via blood and lymph through secondary lymphoid organs using a set of receptors that includes selectin CD62L, chemokine receptor CCR7, and the α_L_β_2_ integrin (CD11a/CD18; aka leukocyte functional antigen 1 [LFA-1]). ^4^ Encounter with the antigen presented in the MHC class I complex on activated antigen-presenting cells (APC) within secondary lymphoid organs stops random migration of naïve T cells and leads to their activation, expansion, and differentiation into cytotoxic effector CD8 T cells (CTLs). Simultaneous upregulation of adhesion molecules, such as β1-integrin (CD29), and inflammatory chemokine receptors, such as CXCR3, licenses CTLs to enter various peripheral organs. ^4^ There, CTLs provide efficient protection against intracellular pathogens through production of the effector molecules granzyme-B and perforin, cytokines interferon-γ (IFNγ) and tumor necrosis factor-α (TNFα). ^2^ After successful elimination of the pathogen, some of the CD8 T cells will become memory cells, which are able to provide a rapid and effective recall response after pathogen re-encounter. ^5^

The generation of a large number of antigen-specific CD8 T cells *ex vivo* enables an elegant way to harness the CTLs to fight tumors and viral infections by adoptive T cell therapy (ATCT). ATCT involves the collection, *ex vivo* expansion, and eventually genetic manipulation of antigen-specific T cells before reinfusion into patients. ^6–8^ ATCT therapies were initially developed to target tumors and have been expanded to tumor-infiltrating lymphocytes, engineered T cell receptor (TCR)-T cells, and chimeric antigen receptor (CAR)-T cells. ^7–9^ ATCT therapies with virus-specific T cells were employed also in immunosuppressed persons and transplant patients for the treatment of viral infections, including those with human cytomegalovirus (HCMV), Epstein-Barr virus (EBV), and BK virus (Human polyomavirus 1). ^6,10^ Among those, HCMV reactivation from lifelong subclinical latent infection is one of the main infectious complications in transplant recipients, reaching incidence of up to 70%. ^11^ Currently, several antiviral drugs are available for the prevention and treatment of HCMV infection. ^12^ However, antiviral therapy is associated with significant toxicity and the emergence of drug-resistant viral strains. ^13^ Hence, the reconstitution of the antiviral immunity by adoptive transfer of donor-derived virus-specific T cells (VSTs) is an attractive alternative treatment strategy in hematopoietic stem cell transplant patients and is also considered for solid organ transplant recipients. ^11–17^

VSTs are generated *ex vivo* from virus-experienced autologous or allogeneic sources, taking care to decrease production time and optimize function after transfer to the patients. ^13,18^ One strategy for ATCT generation for the treatment of viral diseases is to identify and select VST from the donor blood directly using specific peptide-MHC class I complexes ^19,20^ or cytokine capture systems. ^17,21–24^ The second strategy relies on the selective VST expansion after *ex vivo* stimulation of donor leukocytes with antigen-presenting cells presenting viral peptides or proteins. ^16,25–28^ *Ex vivo* expansion protocols are also the basis for genetic engineering approaches, which are particularly interesting as they hold promise to develop VSTs resistant to immunosuppressive drugs, ^29–31^ or to bypass MHC-restriction by producing virus-specific TCR-transgenic T cells ^32^ and CAR-T cells. ^33,34^ At the same time, *ex vivo* expansion strongly affects CD8 T cell phenotype. According to the expression of T cell differentiation markers such as CD45RO, CD45RA, CD27, and CD28, expanded VSTs represent a mixture of cells with effector memory (Tem), central memory (Tcm), CD8 effector memory cell re-expressing CD45RA (TEMRA), and effector phenotypes that varies between the donors. ^16,35,36^ Simultaneously, the expansion affects the expression of homing molecules such as chemokine receptors, integrins, and selectins, affecting expanded T cell circulation within the blood, homing into tissues, and migration to target cells. ^36,37^ However, limited data are available about the circulation and homing behavior of *ex vivo* expanded T cells after intravenous application. A better understanding of *ex vivo* expanded VSTs migration properties could provide important insights in why they do not manage to clear viral infection in some patients. ^16,27^

Here, we describe a simple pre-clinical model that could serve as an experimental system for the comparison of divergent VST therapies. To mimic cytomegalovirus infection in severely immunosuppressed patients, we infected mice deficient for T, B, and NK cells intraperitoneally with different recombinant murine cytomegalovirus (MCMV) strains. These mice could be protected from disease development by the intravenous transfer of T cells expressing recombinant TCR specific for a virus-encoded peptide. As the *ex vivo* expansion of VSTs induced an effector T cell phenotype and led to changes in the integrin and chemokine receptor expression profile, we used CRISPR/Cas9-mediated gene editing to evaluate the role of integrin beta 1 (*Itgb1*; CD29), chemokine receptor CXCR3, and checkpoint molecules PD1 and CLTA4 on the homing and function of *ex vivo* expanded VSTs. Our data indicate that CD29 has an important role in the entry of VSTs with effector T cell phenotype into infected organs. Furthermore, a massive reduction of CD29-deficient VST cell number in blood and spleen suggests that the expansion of pre-activated, adoptively transferred anti-virus effector CD8 T cells primarily occurs in the infected tissues rather than in lymphoid organs. In contrast, the lack of CXCR3 did neither impair entry of adoptively transferred cells into the peripheral tissues nor their localization within tissues and killing of infected cells. The adoptively transferred OT-I cells expressed an array of chemokine receptors, including CCR5 and CXCR6, that seem to contribute significantly to their homing and protective functions. Interestingly, encounters with MCMV-infected cells upregulated checkpoint molecule expression on adoptively transferred OT-I cells. However, transfer of *ex vivo* expanded *Pdcd1*^-/-^ and *Ctla4*^-/-^ OT-I cells provided the same level of protection against MCMV as the transfer of wild-type OT-I cells, arguing against quick exhaustion of *ex vivo* expanded VSTs after transfer.

## 2 Material and methods

### 2.1 Experimental animals

All MCMV infection experiments were done on *Rag2*/IL-2-receptor common gamma chain (Il2rg) double knockout mice (B6-Rag2^tm1Fwa^II2rg^tm1Wjl^ mice; *Rag2*^-/-^*γC*^-/-^) that are deficient for T cells, B cells, and NK cells or B6.Cg-Rag2^tm^^1.1Cgn^/J mice (*Rag2*^-/-^), which are deficient for T and B cells. For additional depletion of NK cells, *Rag2*^−/−^ mice were injected intraperitoneally with 300Lμg NK1.1 antibody (clone PK136, homemade) per animal 4 hr before and every 2 days after infection. As T cell donors, we used C57BL/6-Tg(TcraTcrb)1100Mjb Ptprca-Pepcb/Boy mice (OT-I mice), or F1 generation of pups from intercross of C57BL/6-Tg(TcraTcrb)1100Mjb Ptprca-Pepcb/Boy with either C57BL/6-Tg(CAG-EGFP)131Osb/LeySopJ (GFP^+^OT-I mice) or B6.129(ICR)-Tg(CAG-ECFP)CK6Nagy/J (CFP^+^OT-I mice) strains. *Rag2*^-/-^*γC*^-/-^ were purchased from Taconic Biosciences and all other mouse strains were bred at the central animal facility at Hannover Medical School. All mice were maintained under specific pathogen-free conditions. For experiments, we used female or male mice at the age of 7-30 weeks.

All animal procedures in this study were approved by the Lower Saxony State Office for Consumer Protection and Food Safety (LAVES; TVV 17/2737) and were done following the European and national regulations for animal experimentation, including the European Directive 2010/63/EU and the Animal Welfare Acts in Germany.

### 2.2 Viruses and infections

The MCMV-2D and -3D reporter virus strains, derived from the Smith strain (pSM3fr) as a bacterial artificial chromosome-cloned MCMV progeny, were produced and titrated *in vitro* as described previously. ^38^ Briefly, both mutant strains encode Gaussia luciferase and mCherry inserted into the MCMV m157 open reading frame. MCMV-3D additionally encodes the peptide SIINFEKL in the m164 open reading frame. For infection, 10^6^ PFU of the virus stocks simultaneously titrated by plaque assay on primary murine embryonic fibroblasts (MEF) were diluted in 150 µl phosphate-buffered saline (PBS) and intraperitoneally injected into mice. The time point of injection was defined as the beginning of the infection.

### 2.3 Isolation and purification of CD8 T cells

Cells were isolated from peripheral lymph nodes and spleens from untreated OT-I, GFP^+^OT-I, or CFP^+^OT-I mice. OT-I CD8 T (OT-I) cells were enriched by negative selection using a CD8a^+^ T cell isolation kit (Miltenyi Biotec), commonly reaching a purity of 90-98% as determined by flow cytometry (data not shown). The cells were used either for *in vitro* culture or for administration into mice. In the latter case, we additionally sorted CD8 T cells expressing high levels of GFP or CFP on a FACSAria^TM^ cell sorter (Becton Dickinson) to ensure easier detection upon transfer to the mice.

### 2.4 Activation and proliferation of CD8 T cells in vitro

The purified OT-I, GFP^+^OT-I, or CFP^+^OT-I T cells were cultured in the T cell medium consisting of RPMI medium (ThermoFisher) supplemented with 10% fetal bovine serum (FCS, Sigma-Aldrich), 100 U/ml penicillin (ThermoFisher), 100 µg/ml streptomycin (ThermoFisher), 2 mM L-glutamine (ThermoFisher), 0.05 µM 2-mercaptoethanol (Sigma-Aldrich) and 100 U/ml human IL-2 (Sigma-Aldrich). After initial activation by plate-bound 0.05 µg of anti-CD3 (homemade, Clone: 17-A2) and 0.1 µg of anti-CD28 (clone: 37.51, eBioscience) for 2 days in a 96-well plate, cells were transferred to uncoated plates and expanded for a maximum of 8 days in above mentioned medium before used for flow cytometry analysis or adoptive transfers. In some experiments, at day 7 of culture cells were re-stimulated with the plate-bound 0.05 µg of anti-CD3 and 0.1 µg of anti-CD28 for 18-20 hours in a 96-well plate and used on day 8 for flow cytometry.

### 2.5 CRISPR/Cas9 mediated gene editing

To disrupt gene function, we used clustered regularly interspaced short palindromic repeats (CRISPR)/CRISPR-associated protein 9 (Cas9) ribonucleoproteins (RNPs) as described before. ^39^ Briefly, individual crRNAs (listed in Supplementary Table 1) were mixed at 1:1 molar ratio with tracrRNA (all from Integated DNA Technologies Inc.; IDT), heated at 95°C for 5 minutes in PCR thermocycler and slowly cooled down to room temperature. We used negative control crRNA (IDT) without specificity to mouse, rat, or human genome as a control. Afterward, 210 pmol of each of crRNA:tracrRNA duplexes targeting the same gene were pooled and mixed with Cas9 from *S. pyogenes* (SpCas9, IDT) at 3:1 molar ratio and left at room temperature for 10-20 minutes to form CRISPR/Cas9 RNPs. Just before nucleofection 70 pmol of electroporation enhancer (100 nucleotide long single-stranded DNA molecules without any similarity to human, mouse, or rat genome; IDT) was added to the mix.

For nucleofection, up to 2x10^6^ of OT-I cells, pre-activated with plate-bound anti-CD3 and-CD28 for 2 days as described above, were nucleofected using P3 Primary cell 4D-nucleofection kit (Lonza) and DN100 pulse on 4D nucleofector X unit (Lonza). After nucleofection, the cells were transferred into 1.5 ml of T cell medium and further cultured as described above. After 2 days, nucleofection was repeated to increase gene deletion efficacy. One day before cell transfer, we additionally sorted knockout cells using FACS Aria after labeling the remaining WT cells using appropriate antibodies as described in subchapter 2.11.

#### CRISPR/Cas9-mediated

To determine CRISPR/Cas9-induced gene deletion scores for Cas9 RNPs targeting *Ctla4* and *Pdcd1*, we extracted DNA from untouched and nucleofected cells using QIAamp DNA Mini Kit (Qiagen). Next, we amplified the targeted gene regions by PCR (NEB Next High-Fidelity 2xPCR Master Mix; New England Biolabs) using primers (Supplementary Table 2) and purified the products using QIAquick PCR Purification Kit (Qiagen). PCR-products were Sanger sequenced and knockout scores were determined using the ICE analysis tool (https://ice. synthego.com/, Synthego). ^40^

### 2.6 Adoptive transfer of CD8 T cells

In all the adoptive T cell transfer experiments, CD8+ T cells were diluted in 100 µl sterile PBS and transferred to MCMV-infected *Rag2*^-/-^*γC*^-/-^ or *Rag2*^-/-^ mice by intravenous injection at one day post-infection (dpi). In initial experiments, MCMV-3D or MCMV-2D-infected *Rag2*^-/-^*γC*^-/-^ mice received 1×10^5^ naïve GFP^+^OT-I T cells. To study the role of integrin beta 1 in the migration of activated virus-specific T cells, MCMV-3D-infected *Rag2*^-/-^*γC*^-/-^ mice received a mixture of *in vitro* activated 1×10^4^ *Itgb1*^+/+^ CFP^+^OT-I T cells and 1×10^4^ *Itgb1*^-/-^ GFP^+^ OT-I T cells. To examine the role of CXCR3, CTLA4, and PD-1 in migration and/or protective function of antiviral CD8 T cells, MCMV-3D-infected *Rag2*^-/-^ mice received 7.5×10^4^ GFP-OT-I T cells or the same number of *Cxcr3*^-/-^*, Ctla4*^-/-^, or *Pdcd1*^-/-^ GFP-OT-I T cells.

### 2.7 Sample collection

At 2-12 dpi mice were sacrificed using CO_2_ anesthesia and cervical dislocation. In the experiments with Rag2^-/-^γc^-/-^ mice, blood was withdrawn from the hearts of the mice receiving adoptive T cell transfer. Afterward, all mice were perfused with 20 ml of ice-cold PBS transcardially to remove blood cells before organ collection. Next, the spleen, liver, lung, and salivary gland (SVG), were harvested for further analysis. The following portions of the organs were stored in cold 500 µl DMEM containing 10% FCS, 100 U/ml penicillin, and 100 µg/ml streptomycin for luciferase assay: pieces from the same anatomical parts of spleens (weight range 5-15 mg), livers (weight range 190-220 mg), left salivary glands, and left lung lobes. Small pieces (weight range 5-10 mg) from the same anatomical parts of spleens were collected and stored in cold FACS buffer (PBS supplemented with 3% FCS) for cell isolation. The rest of the organs was fixed in a cold 2% paraformaldehyde (PFA) solution containing 30% sucrose and used for histological analysis.

The Rag2^-/-^ mice received intravenously 10 µg of Thy1.2-PerCP-Cy5.5 or CD45.2-PerCP antibody 5 minutes before organ collection to stain leukocytes within or close to the blood vessels. In those experiments, spleens, livers, and lungs were collected and separated for luciferase assay, flow cytometry, and histology as described above.

### 2.8 Luciferase assay

To monitor the spread of the virus, organ parts collected as described above were homogenized by TissueLyser II (Qiagen, 25Hz, 4 min) and centrifuged to remove debris. For luciferase assay, 180 µl of the 1:10 diluted supernatants were mixed with 20 µl of the 1 µg/ml Coelenterazine substrate (Invitrogen), and luciferase activity within the mixture was measured in duplicates with Lumat LB 9507 (Berthold Tech) for 10 s. To normalize the values across the experiments, we divided the measured luciferase activity with the weight of the tissue used for homogenization to express values as relative light units (RLU) per mg of tissue.

### 2.9 Histology

After fixation in a 2% PFA solution containing 30% sucrose at 4°C for 24 hours, the organs were embedded in the O.C.T. Compound (Tissue-Tek, Sakura) and sectioned using CM3050 S Cryostat (Leica). If antibody staining was performed, cryosections (7-8 µm thick) were blocked with 10% rat serum and Fc receptor block (Clone: 2.4G2, homemade) solution before staining with APC-labeled anti-CD45 (Supplementary Table 3). In all sections, nuclei were labeled with 4’,6-Diamidino-2-phenylindole dihydrochloride (DAPI; Sigma-Aldrich) or TO-PRO-3^TM^ (Invitrogen). The immunofluorescent photomicrographs were acquired with the Axiovert 200M inverted microscope (Carl Zeiss; PlanApochromat objectives 10x/0.45 and 20x/0.75 (magnification/numerical aperture) and AxioCam MRm camera) or with AxioScan Z1 (Carl Zeiss; Plan-Apochromat objective: 10x/0.45 M27; camera: Axiocam 506 mono), contrast adjusted, and analyzed using AxioVision Rel. 4.8 or ZEN blue 2.3 (both Carl Zeiss).

### 2.10 Isolation of cells from organs for flow cytometry

Collected lung and liver lobes were cut into small pieces and digested for 45 minutes at 37°C in digestion media containing (i) RPMI supplemented with 5% FCS, 2.5% Hepes, 0.025 mg/ml DNAse I (#11284932001, Roche), and 0.5 mg/ml Collagenase D (#11088866001 Roche) for lungs or (ii) RPMI supplemented with 10% FCS, 5% Hepes, 0.025 mg/ml DNAse I (#11284932001, Roche), and 0.5 mg/ml Collagenase D (#11088866001 Roche) for liver. After digestion, single cells were separated from the organs by smashing the remaining tissue parts through 70 µm cell strainers. Single-cell suspensions from the spleen were obtained by smashing the organs through 70 µm cell strainers. Before cell counting, single-cell suspensions of all organs were subjected to hypertonic red blood cell lysis.

### 2.11 Flow cytometry

Flow cytometry was used to analyze T cell purity, activation status, protein expression after genome editing, and/or phenotype and expansion after adoptive transfer. For flow cytometric analyses, we used antibodies and other reagents listed in Supplementary Table 3. Briefly, the staining protocol included initial blocking of non-specific antibody binding by incubating samples with 10% rat serum for 10 minutes at 4°C. Next, primary antibodies were added, and cells were stained at 37°C with mixtures containing chemokine receptor antibodies or at 4°C with all other antibody mixtures. After washing, cells were either acquired or stained with streptavidin for an additional 20 minutes at 4°C, washed, and then used for acquisition. The acquisition was done on BD™ LSR II or Symphony A1 cytometers (BD Biosciences, San Jose, CA), and data were analyzed with FlowJo 10.4.1 (Tree Star Inc.).

### 2.12 Statistics

For statistical analyses, we used Prism 7 or 8 (GraphPad) using tests as listed under each figure.

## 3 Results

### 3.1 Intravenous transfer of antigen-specific CD8 T cells protects Rag2 ^-/-^γC^-/-^ mice from MCMV infection

To establish a model of the cytomegalovirus infection in severely immunosuppressed transplant patients, we initially used *Rag2*^-/-^*γC*^-/-^ mice, which are deficient for T cells, B cells, and NK cells. For infection, we have chosen recombinant MCMV-3D. This MCMV strain encodes for the fluorescent protein mCherry and a secretable Gaussia luciferase, allowing elegant tracking of the infected cells and the kinetic analyses of virus spread in various organs. ^38,41^ Additionally, MCMV-3D encodes for the SIINFEKL peptide originating from chicken ovalbumin (residues 257-264). Since SIINFKL is presented on H-2k^b^ MHC-I molecules by infected cells and is recognized by a transgenic T cell receptor carried by OT-I cells, CD8 T cells isolated from OT-I mice served as a convenient cell source for ATCT. However, the MCMV-3D virus strain also carries a frameshift mutation in the MCMV open reading frame *m131/m129* that encodes for the MCMV-encoded chemokine 2 (MCK2). ^42^ As a result, MCK2-deficient MCMV-3D is impaired in entry into macrophages and fails to disseminate efficiently to the salivary gland upon intranasal infection. ^42–44^ To assure that MCMV-3D dissemination upon systemic infection is unaffected by the lack of MCK2, we initially infected *Rag2*^-/-^*γC*^-/-^ mice intraperitoneally with 10^6^ PFU of MCMV-3D (Fig. 1A). Already at 2 dpi the virally encoded Gaussia luciferase could be detected in the spleen and at 4 dpi in liver, lungs, and salivary glands (Fig. 1B). The amount of virus, measured as Gaussia luciferase activity per mg of tissue weight, steadily increased in all organs except in the spleen, where it remained relatively constant. Eventually, the MCMV overwhelmed the protection, exhausting the mice that had to be euthanized at 10-12 dpi as they lost up to 10% of their weight. Altogether, these initial experiments confirmed the importance of the adaptive immune system and NK cells in the control of MCMV infection. ^45–47^

**Figure 1.**
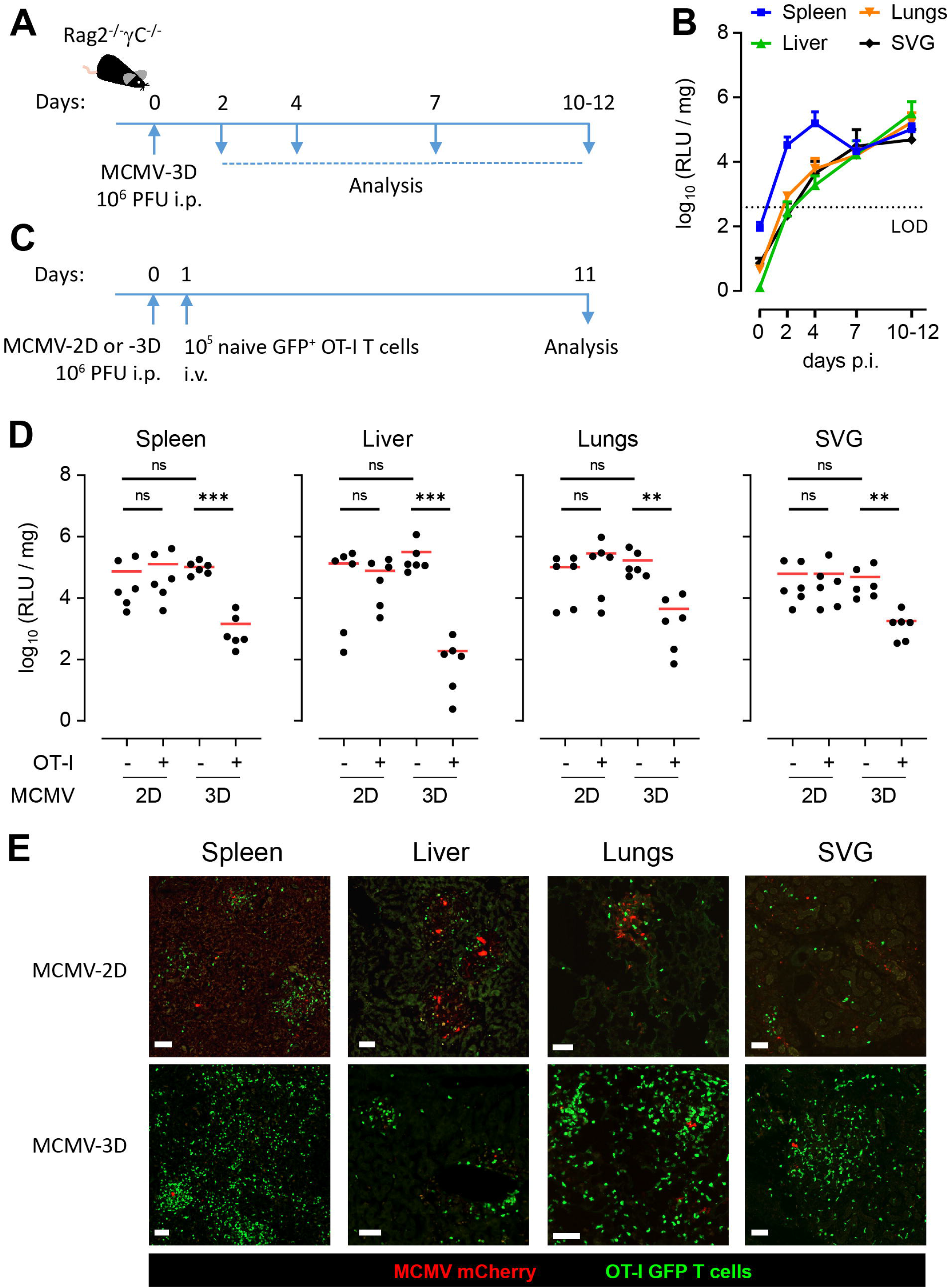
Antigen-specific T cells protect immunodeficient *Rag2*^-/-^*γC*^-/-^ mice from MCMV infection. (**A**) Experiment scheme for infection analysis. (**B**) MCMV-3D infection in indicated organs (SVG – salivary gland) measured as Gaussia luciferase activity per mg of tissue at indicated days post infection (dpi). Pooled data from 5 experiments, shown as mean ± SD from 3 (0 dpi) or 6 (2, 4, 7, 10-12 dpi) mice per group. (**C**) Experimental scheme for adoptive T cell therapy. MCMV-2D and -3D encode for mCherry and secreted Gaussia luciferase. MCMV-3D additionally encodes for SIINFEKL peptide recognized in context of H-2Kb molecules by OT-I CD8 T cells. (**D**) Gaussia luciferase activity at 12 dpi with MCMV-2D or -3D in indicated organs of mice that did (+) or did not (-) intravenously receive 10^5^ OT-I T cells one day after infection. Pooled data from 4 experiments, shown as individual mice (dots) and group mean (line). Data from MCMV-3D infected Rag2^-/-^γC^-/-^ mice without OT-I T cell transfer are identical to those shown in Figure 1B. Statistical analysis was done on log-transformed values using ordinary or Welch’s ANOVA followed by Dunnett’s T3 multiple comparisons test. **p < 0.01, ***p < 0.001. (**E**) Representative photomicrographs of indicated organs reveal swarming of transferred OT-I T cells around cells infected with MCMV-3D but not -2D at 12 dpi. Scale bar – 50 µm.

To validate that OT-I cells provide antigen-specific protection against MCMV-3D infection, we intravenously transferred 10^5^ isolated GFP^+^OT-I T cells in mice that were infected with MCMV-3D or MCMV-2D (Fig. 1C). MCMV-2D is a virus strain that shares all genetic modifications with MCMV-3D except that it does not encode for the SIINFEKL peptide. ^38,41^ As expected, the adoptive transfer of OT-I cells significantly reduced the viral load in all investigated organs of MCMV-3D- but not MCMV-2D-infected mice (Fig. 1D). Interestingly, OT-I cells also controlled viral infections in salivary glands (Fig. 1D), whose protection has been suggested to primarily rely on CD4 T cells. ^48^ Histological analysis of organs from MCMV-2D and -3D infected animals at 12 dpi indicated that GFP^+^OT-I cells accumulated around both MCMV-2D- and MCMV-3D infected cells (Fig. 1E). In MCMV-2D-infected animals, however, mCherry^+^ MCMV-infected cells were readily detectable and organized in foci (Fig. 1E), further confirming uncontrolled viral spread within the tissue. In contrast, in organs from MCMV-3D infected animals GFP^+^OT-I cell infiltrates were much denser and often did not contain mCherry^+^ MCMV-infected cells, indicating that those were eliminated and no longer detectable (Fig. 1E).

Together, these data indicate that intravenous transfer of OT-I cells into MCMV-3D infected *Rag2*^-/-^ *γC*^-/-^ mice represents a simple pre-clinical model that could serve as an experimental system for the comparison of divergent VST therapies.

### 3.2 Ex vivo OT-I CD8 T cell expansion changes the expression repertoire of homing molecules

Many protocols for the generation of VSTs for ATCT involve *ex vivo* cell expansion, especially those involving genetic manipulation of the cells. ^18^ *Ex vivo* expansion, however, might affect the phenotype of VST. To study such an effect in our model, we activated isolated naïve OT-I cells *in vitro* by stimulation with plate-bound anti-CD3 and anti-CD28 antibodies in the presence of IL-2. After 2 days, cells were removed from the plates and further expanded for 5 days in a medium containing IL-2. At day 7 of culture, cells had increased size, upregulated CD25 (IL-2Rα), retained CD62L expression, but did not upregulate CD44 (Fig. 2A and Supplementary Fig. 1A). As expected, *ex vivo* expansion affected the expression of homing molecules. Compared to freshly isolated cells, *ex vivo* expanded OT-I cells at day 7 of the culture showed higher expression of integrin αv (CD51) and integrin β1 (CD29) but lower expression of integrin β7 (Fig. 2B). At the same time, integrin β2 (CD18) remained highly expressed (Fig. 2B). Activated OT-I cells also downregulated chemokine receptor CCR7 and strongly upregulated expression of CCR5, CXCR3, and CXCR6 (Fig. 2C). CCR1, CCR2, CCR4, CCR6, CCR9, CXCR5, and CX3CR1 were expressed at low levels or not expressed at all in either naive or expanded OT-I cells (Supplementary Fig. 1B). Together, these data indicate that *ex vivo* expansion changes homing properties of OT-I cells, reducing expression of homing molecules that would allow T cell entry into secondary lymphoid organs (CCR7 and integrin β7) and upregulating receptors crucial for their entry into peripheral tissues (CCR5, CXCR3, and integrin β1).

**Figure 2.**
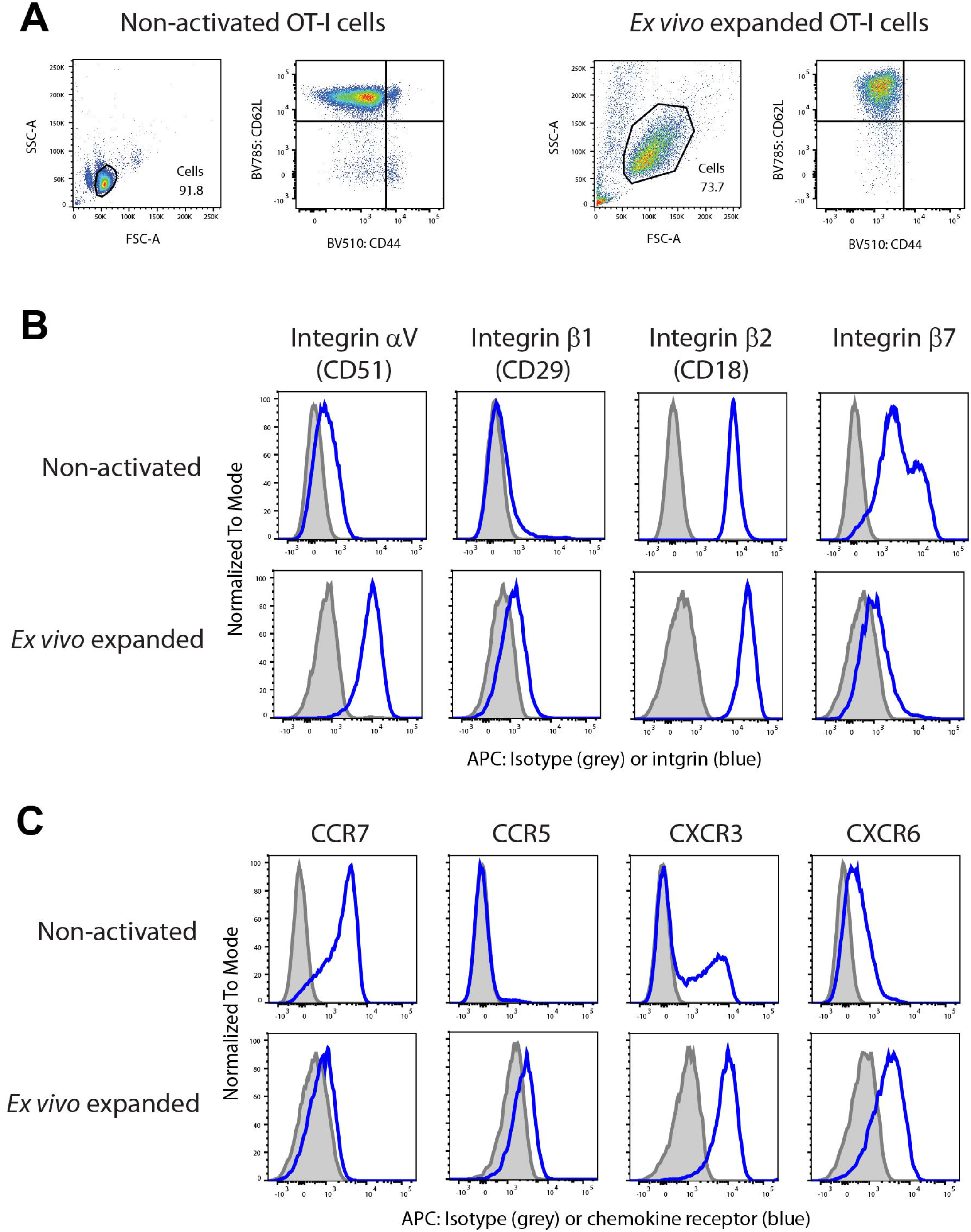
*Ex vivo* expansion alters expression of homing molecules on OT-I CD8 T cells. (**A**) Representative pseudocolor plots showing the forward and side scatter and the expression of CD62L and CD44 of CD8 OT-I T cells before (left) and after 7 days of *ex vivo* expansion (right). (**B-C**) Expression of (**B**) integrin subunits and (**C**) chemokine receptors on CD8 OT-I T cells before (non-activated) and after 7 days of *ex vivo* expansion: blue lines, antibody staining; gray shaded areas, isotype control. (**A-C**) Representative plots and histograms from 2-3 experiments.

### 3.3 Integrin-β1 is required for the in vivo proliferation of intravenously transferred VSTs

Marked upregulation of homing molecules important for entry into peripheral tissues indicated that adoptively transferred *ex vivo* expanded VSTs might provide protection without additional proliferation *in vivo* within secondary lymphoid organs. To enter infected peripheral organs from blood, T cells use integrin α4β1 [very late antigen 4 (VLA4)] for the firm adhesion on vascular cell adhesion molecule-1 (VCAM-1, CD106) expressed on endothelial beds at sites of inflammation. ^49,50^ To investigate the function of adoptively transferred VSTs in a situation where they are prevented from entering infected organs, we knocked out *Itgb1* with nucleofection delivered CRISPR/Cas9-RNP as described earlier (Fig. 3A). ^39^ We co-transferred equal numbers of *ex vivo* expanded CD29^+^CFP^+^ and CD29^-^GFP^+^ OT-I cells one day after MCMV-3D infection and examined the organs at 11 dpi (Fig. 3B). As expected, we observed reduced numbers of CD29^-^GFP^+^ OT-I cells in lungs and liver (Fig. 3C). Surprisingly, the numbers of CD29^-^GFP^+^ OT-I cells were also massively reduced in blood and spleen and on average only reached 4% of the CD29^+^CFP^+^ OT-I cell counts (Fig. 3D and Supplementary Fig. 2). These findings suggest that the entry of transferred *ex vivo* expanded VSTs into peripheral infected organs is crucial for their proliferation after transfer and that this proliferation primarily takes place in the infected tissues.

**Figure 3.**
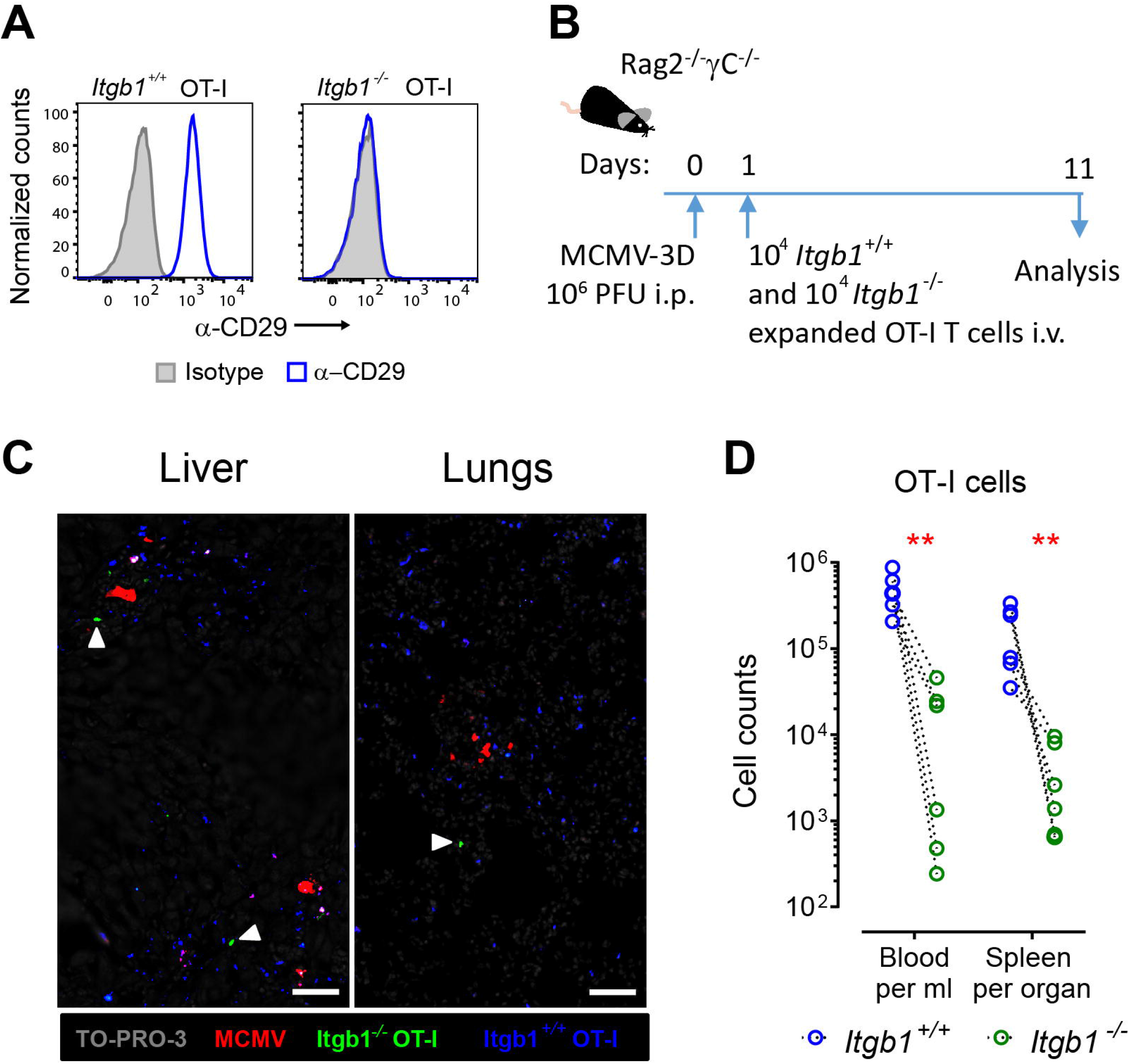
Integrin-β1 (CD29) mediates the proliferation of adoptively transferred *ex vivo* expanded OT-I cells. (**A**) Representative histograms showing CD29 surface expression after CRISPR/Cas9-mediated deletion using a negative control gRNA or three different gRNA targeting the *Itgb1* (N = 3 experiments). (**B**) Experimental scheme of ATCT. (**C**) Representative photomicrographs of liver and lung sections examined for the presence of *Itgb1*^+/+^ CFP^+^ and *Itgb1*^-/-^ GFP^+^ OT-I cells at 11 dpi (N = 6). Scale bar: 50 µm. Arrowheads mark rare *Itgb1*^-/-^ GFP^+^ OT-I cells. (**D**) Numbers of CD29^+^CFP^+^ and CD29^-^ GFP^+^ OT-I cells at 11 dpi in blood and spleen of mice that received 10^4^ of each cell population one day before MCMV-3D intraperitoneal infection. Pooled data from two experiments, shown as individual mice (dots). Ratio paired t-test. ** p < 0.01.

### 3.4 CXCR3 is not required for homing of ex vivo expanded VSTs into infected organs

In a three-step cascade of T cell immigration from blood into inflamed organs, integrin adhesiveness to their substrates is activated through signals transduced from chemokine receptors bound to chemokines displayed on the surface of the epithelium or released from the tissue. Upon entry into the tissue, chemokine receptors are crucial to guide VSTs to close proximity of infected cells within a complex tissue environment. Among the chemokine receptors upregulated on *ex vivo* expanded VSTs, we decided to investigate the role of CXCR3 in that process. This chemokine receptor has been implicated in CD8 T cell interactions with APCs in secondary lymphoid organs, entry of effector cells into infected organs and the precise positioning of T cells within tissues. ^4,51–53^ Successful CRISPR/Cas9-mediated gene editing enabled us to investigate the role of CXCR3 in the migration of adoptively transferred *ex vivo* expanded OT-I cells (Fig. 4A and B). In these experiments we switched to *Rag2*^-/-^ mice as recipients. To assure that NK cells were also depleted, we administered intraperitoneally 300Lμg NK1.1 antibody (clone PK136) per mouse 4 hr before and every second day after infection. Flow cytometric analysis confirmed complete NK depletion (data not shown). Interestingly, we found that CXCR3^-^ OT-I cells protected mice from the MCMV-3D infection comparably as their wild-type counterparts (Fig. 4C). Quantification of transferred cells within the infected organs indicated that CXCR3^+^ and CXCR3^-^ OT-I cells accumulated equally well within spleens, livers, and lungs (Fig. 4D and Supplementary Fig. 3). Histological analysis of livers and lungs at 8 dpi indicated that both CXCR3^+^ and CXCR3^-^ OT-I cells accumulated within nodular inflammatory foci composed of CD45^+^ leukocytes surrounding mCherry^+^ MCMV-3D infected cells (Fig. 4E). Interestingly, in spleens of the same animals we detected mCherry mainly in small cell fragments, suggesting that in this organ both CXCR3^+^ and CXCR3^-^ OT-I cells already eliminated the majority of infected cells. To validate which other chemokine receptors mediate the migration of adoptively transferred cells within the infected organs, we additionally profiled chemokine receptor expression on CXCR3^+^ OT-I cells isolated at 8 dpi from MCMV-3D infected organs. Our analysis indicated that these cells, besides CXCR3, express CCR5 and CXCR6 (Supplementary Fig. 4), and thus maintain their chemokine receptor expression profile from *ex vivo* expansion (Fig. 2 and Supplementary Fig. 1). Together, these data indicate that tissue homing and migration of adoptively transferred VSTs does not rely on CXCR3.

**Figure 4.**
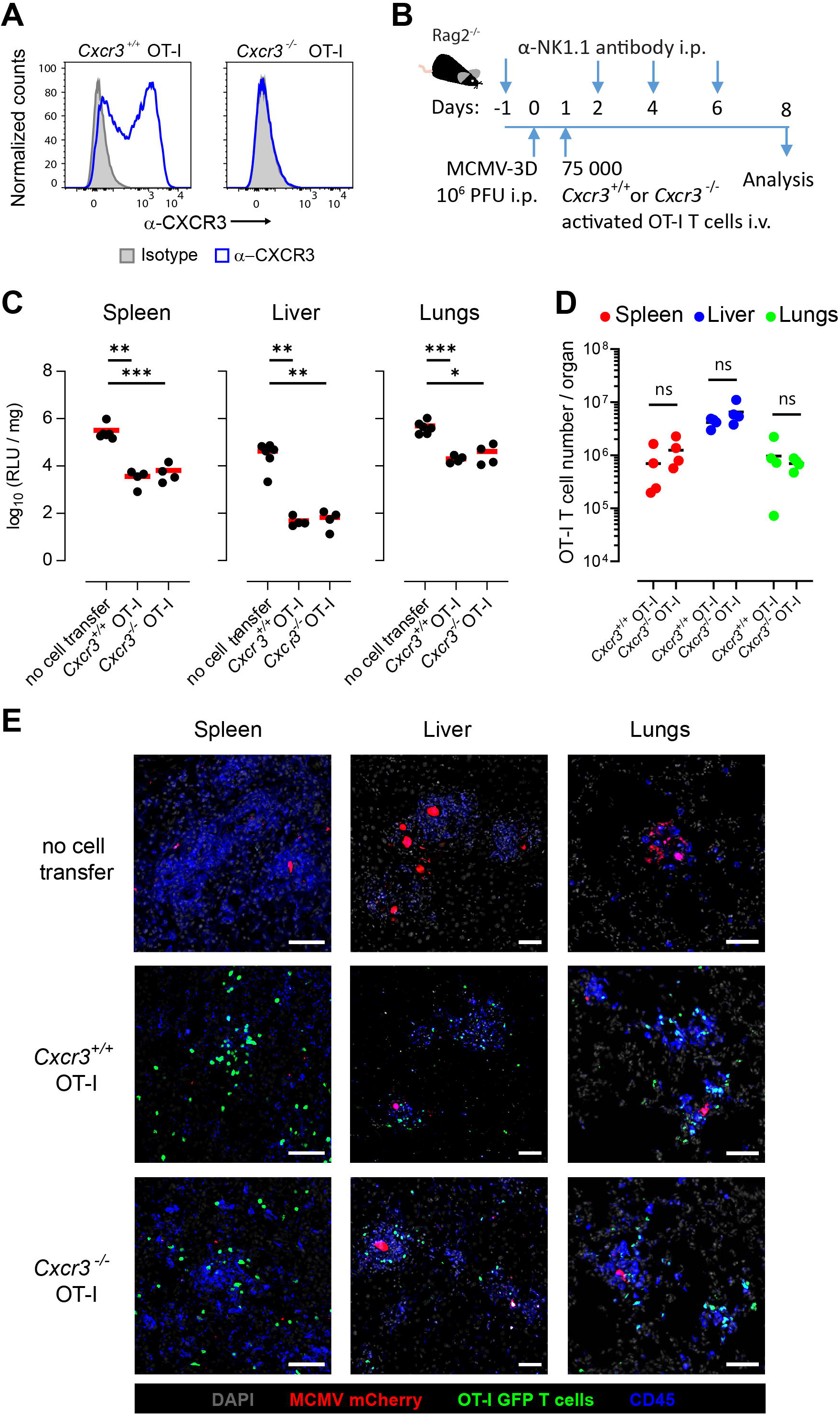
CXCR3 is not required for entry of *in vitro* expanded antiviral T cells into peripheral organs. (**A**) Representative histograms showing CXCR3 surface expression after CRISPR/Cas9-mediated deletion using a negative control gRNA or three different gRNA targeting the *Cxcr3* gene (N = 5 experiments). (**B**) Experimental scheme of ATCT. (**C**) Gaussia luciferase activity at 8 dpi within spleens, livers, or lungs of mice that intravenously received 7.5 x 10^4^ *Cxcr3*^+/+^ or Cxcr3^-/-^ GFP^+^OT-I CD8 T cells or were left untreated one day after infection. Pooled data from 4 experiments, shown as individual mice (dots) and group mean (line). Statistical analysis was done on log-transformed values using ordinary or Welch’s ANOVA followed by Dunnett’s T3 multiple comparisons test. ** p < 0.01, *** p < 0.001. (**D**) Numbers of *Cxcr3*^+/+^ GFP^+^ and *Cxcr3*^-/-^ GFP^+^ OT-I cells at 8 dpi in indicated organs. Pooled data from four experiments, shown as individual mice (dots) and group means (lines). Unpaired t-test with Welch’s correction. ns - p > 0.05. (**E**) Representative immunofluorescence photomicrographs of organs showing positioning of transferred *Cxcr3*^+/+^ GFP^+^ or Cxcr3^-/-^ GFP^+^ OT-I CD8 T cells around MCMV-3D infected cells at 8 dpi. Scale bar – 50 µm.

### 3.5 Checkpoint molecules CTLA4 and PD1 do not limit anti-viral VST response

Our results indicated that adoptively transferred VSTs enter the infected tissue where they provide immediate protection. During flow cytometric analysis of adoptively transferred VSTs, we noticed that they acquire expression of checkpoint molecule PD-1 but not CTLA4 (Fig. 5A). Restimulation of *ex vivo* expanded OT-I cells with plate-bound anti-CD3 and anti-CD28 antibodies confirmed that their activation resulted in increased expression of both PD-1 and CTLA4 (Fig. 5B). Recent data indicated that therapeutic blockade of checkpoint molecules can augment T cell effector functions not only against tumors but also during acute viral infections. ^54^ Thus, we investigated whether the CRISPR/Cas9 RNP-mediated deletion of the checkpoint molecules PD1 or CTLA-4 in *ex vivo* expanded OT1 T cells might also have a beneficial role in anti-MCMV immune responses. Since the low level of CTLA4 expression on the cell surface prevented clear distinction of *Ctla4*^-/-^ and wild-type cells, we evaluated the success of modifications using gene editing efficiency analysis. ^40^ Analysis of gene loci targeted by single CRISPR/Cas9 RNPs still contained up to 30% of intact sequences (Fig. 5C). Thus, we targeted *Pdcd1* and *Ctla4* using a combination of 3 and 2 CRISPR/Cas9 RNPs, respectively, to assure that the genes will remain intact in <5% of cells. The success of *Pdcd1* gene editing was confirmed by flow cytometry (Supplementary Fig. 5).

**Figure 5.**
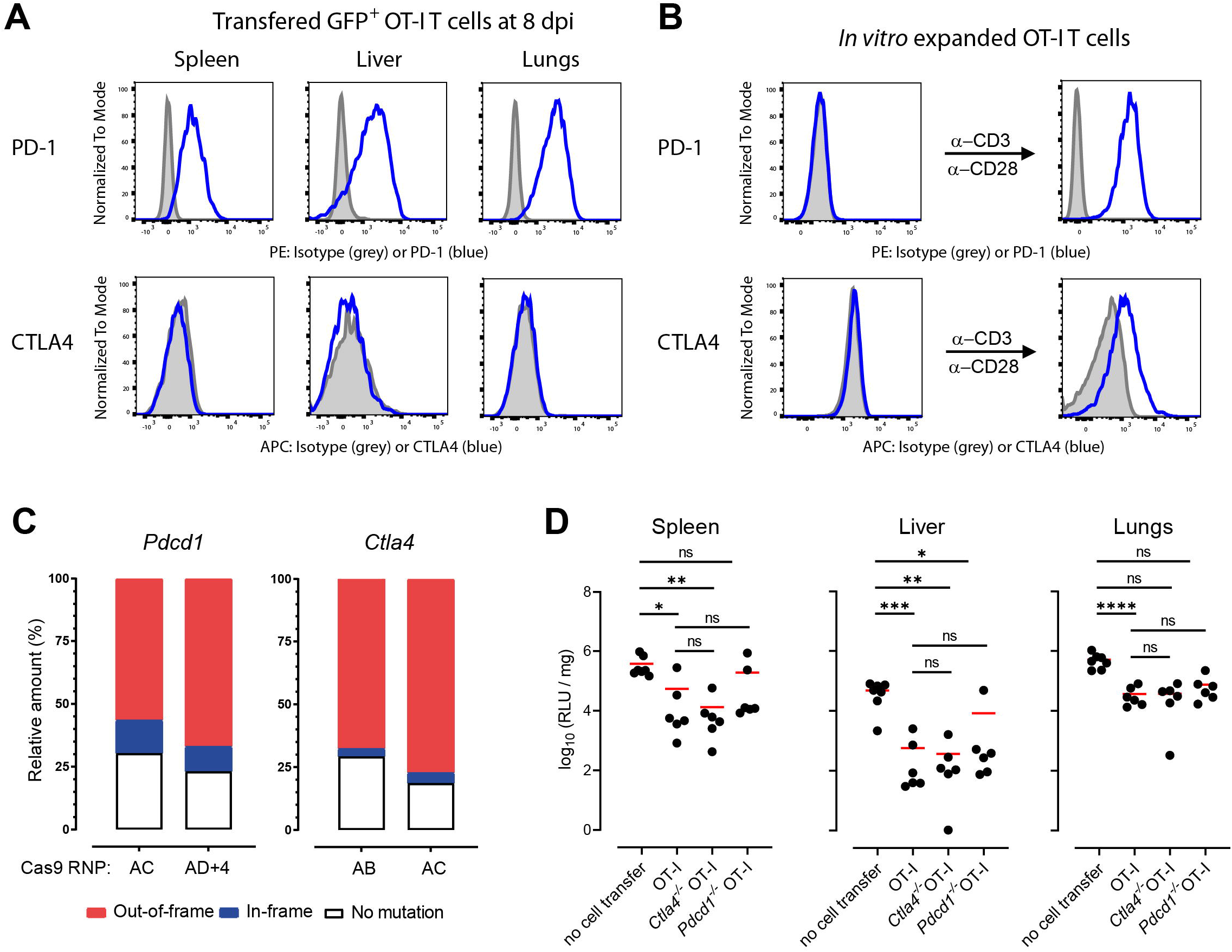
Checkpoint molecules PD-1 and CTLA4 have no effect on CD8 T cell-mediated anti-MCMV immune response. Representative histograms showing PD-1 and CTLA4 surface expression on (**A**) adoptively transferred GFP^+^OT-I cells within indicated organs from *Rag2*^-/-^ mice at 8 dpi with MCMV-3D (N = 5) and (**B**) *ex vivo* expanded GFP^+^OT-I cells at day 7 of culture (left) or after overnight restimulation with anti-CD3/anti-CD28 antibodies. (**C**) Relative distribution of mutations within *Pdcd1* and *Ctla4* genes achieved with different Cas9 RNPs. (**D**) Gaussia luciferase activity at 8 dpi within spleens, livers, or lungs of mice that intravenously received *Pdcd1*^-/-^, *Ctla4*^-/-^, or control GFP^+^ OT-I CD8 T cells or were left untreated one day after infection. Pooled data from 5 experiments, shown as individual mice (dots) and group mean (line). Statistical analysis was done on log-transformed values using ordinary or Welch’s ANOVA followed by Dunnett’s T3 multiple comparisons test. ns – p > 0.05, * p < 0.05, ** p < 0.01, *** p < 0.001, **** p < 0.001. Six data points from MCMV-3D infected Rag2^-/-^ mice without OT-I CD8 T cell transfer and four from MCMV-3D infected Rag2^-/-^ mice that received control OT-I CD8 T cells are identical to those shown in Figure 4D.

Applying the model described above, we infected NK cell-depleted Rag2^-/-^ mice with 10^6^ PFU MCMV-3D. One day later, some of the mice received OT-I cells or OT-I cells deficient for *Ctla4* or *Pdcd1*. When analyzing in the recipients 7 days later, *ex vivo* OT-I cells deficient for CTLA-4 or PD1 reduced the virus loads in the spleen, liver, and lungs to the same extent as the control OT-I cells (Fig. 5D).

These data indicate that PD1 and CTLA-4 do not substantially limit anti-viral properties of *ex vivo* expanded CD8 T cells.

## 4 Discussion

Numerous clinical studies confirmed the efficacy of adoptively transferred VSTs for viral infection treatment in immunosuppressed transplant patients, including those with cytomegalovirus infection or reactivation. ^6,10–17^ Strategies used to generate VST immunotherapy often use *ex vivo* expansion, especially those involving genetic manipulations of the cells or their generation from third-party donors. ^16,25–30,32–34^ However, correlations between the expanded cell phenotype and clinical efficacy are still lacking. We therefore developed a simple pre-clinical model as an elegant experimental system to investigate the homing properties of adoptively transferred *ex vivo* expanded VSTs. In this model, we could show that anti-CD3/anti-CD28-stimulated OT-I cells depend on CD29 but not CXCR3 to enter MCMV-infected organs. The entry into the peripheral tissues was crucial for their *in vivo* proliferation and protective function. Our data further suggest that CCR5 and/or CXCR6 mediate the accumulation of adoptively transferred *ex vivo* expanded OT-I cells and that their protective function is not regulated by checkpoint molecules CTLA4 and PD-1.

Our data highlight the importance of the homing properties of divergent ATCT preparations for the success of the therapy. Importantly, proliferation and, consequently, the efficacy of adoptively transferred VSTs with effector phenotype in our model was dependent on their ability to enter into the organs and contact infected cells. This finding helps to explain why T cells expanded *ex vivo* with anti-CD3/anti-CD28 antibody stimulation do not efficiently protect against tumors in preclinical models. ^55^ In our model, deletion of CD29 impaired the cells to use the VLA4-VCAM-1 axis to enter infected organs. ^49,50^ However, this axis might not be involved in the migration of adoptively transferred T cells into tumors. As antigen encounter within peripheral tissues seems to be a prerequisite for *in vivo* proliferation of adoptively transferred effector CD8 T cells, ATCT efficacy against tumors could be increased using *ex vivo* expansion protocols that equip cells with the appropriate homing molecules to enter the tumor they target.

In addition to integrins and further adhesion molecules, migration patterns of adoptively transferred T cells are also mediated by the expression of chemokine receptors, which are crucial for guiding immune cells within lymphoid and nonlymphoid tissues. ^56,57^ Here, we focused on the role of CXCR3, an inflammatory chemokine receptor involved in multiple stages of CD8 T cell function, including their priming in secondary lymphoid organs, homing to and migration within the infected tissues, and memory formation. ^52,58^ Interestingly, our data indicate that the lack of this chemokine receptor did not impair adoptively transferred OT-I cells to find and destroy MCMV-infected cells in the spleen, lungs, and liver. At first glance, these findings contradict previous studies suggesting that CXCR3 mediates the recruitment of antigen-specific T cells or their migration in various organs during viral infections, including those with MCMV. ^59–64^ However, those studies employed *Cxcr3*^-/-^ mice or transfer of naïve *Cxcr3*^-/-^ T cells into wild-type animals before infection. Recent data suggest that CXCR3 expression is crucial for CD8 T cell interaction with APCs during priming and differentiation into short-lived effector cells within secondary lymphoid organs. ^65–68^ Hence, impaired differentiation of *Cxcr3*^-/-^ CD8 T cells could also explain their subsequent reduced accumulation in infected organs. In our model, w e disrupted *Cxcr3* after initial T cell activation. Therefore, data presented in this study suggest that CXCR3 is not involved in homing or migration of effector CD8 T cells during viral infection. Our characterization of transferred OT-I cells within the infected organs suggests a role of two other expressed chemokine receptors, CCR5 and/or CXCR6, in organ recruitment of effector VSTs. Indeed, both chemokine receptors have been implicated in the migration of effector CD8 T cells at sites of infection. ^69–73^ Hence, it would be important to address the role of CCR5 and CXCR6 for the migration of adoptively transferred *ex vivo* expanded VSTs in future experiments.

Of note, our expansion protocol with anti-CD3 and anti-CD28 antibody stimulation resulted in OT-I cells with uniform effector T cell phenotype. Interestingly, CD62L expression remained high on the surface of expanded cells, which might boost their efficacy upon adoptive transfer. Recent data from preclinical models suggested that virus-specific CD62L^+^ Tcms provide better control of MCMV infection than the Tems or effector T cells. ^74^ The increased protective effect was attributed to higher Tcms proliferation within the infected organs. ^74^ Interestingly, the efficacy of expanded T cells against tumors also depended on the perseverance of CD62L expression that enabled their activation within tumors. ^37,75^ The adhesive function of CD62L might improve the binding of adoptively transferred T cells with antigen-presenting cells within the tissues, therefore increasing their activation and subsequent proliferation. ^75^

The observed upregulation of the checkpoint molecule PD-1 served as a further piece of evidence for the activation of adoptively transferred OT-I cells in the infected organs. Prolonged antigen exposure in tumors and during chronic infections leads to permanent expression of checkpoint molecules that limit T cell function. Targeting those molecules with monoclonal antibodies to inhibit their function and release T cells from blockade has shown success in clinical trials for cancer treatment and is evaluated for the treatment of chronic infections. ^54,76,77^ The role of checkpoint molecule expression on T cells during acute infections is, however, less well understood. ^54,78,79^ Several studies have suggested that checkpoint molecules limit T cell function also during acute viral infections. ^80–83^ On the other hand, our data are more in line with the studies describing that checkpoint molecules do not directly inhibit antiviral T cell expansion or T cell effector functions in acute disease. ^84–86^

Altogether, these data strongly imply that it would be crucial to address the homing properties of other phenotypes contained in therapeutic products used in clinical studies. ^16,35,36^ These experiments might provide important insights into why ATCTs do not show protective effects in all patients. ^16^ Further, our data imply that future clinical evaluation of ATCT efficacy should include homing molecule expression assessment and that ATCT production strategies should use protocols that equip T cells with homing molecules crucial for their entry into target (peripheral) tissues. In future studies, the present model could serve to validate different anti-viral immunotherapy strategies. Moreover, it could also be adapted for testing various CRISPR-Cas9-induced modifications of virus-specific T cells directed towards the improvement of virus control.

### 4.1 Limitations of the study

Similar to other murine models for immunotherapy of CMV-disease, our model also has potential limitations. ^87^ Apart from strict host-species restrictions, Rag2^-/-^γC^-/-^ and Rag2^-/-^mice, in contrast to transplant patients, were not exposed to immuno-suppressives that could affect the function of the transferred T cells. Moreover, our model involves primary systemic infection rather than virus re-activation.

## Acknowledgments

We would like to thank Ms. Svetlana Piter for providing excellent animal care and Dr. Matthias Ballmaier of the Cell Sorting Core Facility of Hannover Medical School for his superb service.

## Funding

Research in the Förster lab is funded by the German Center for Infection Research TTU 01.938 (grant no 80018019238); by the German Center for Lung Research (grant 82DZL002B1); and by Deutsche Forschungsgemeinschaft (DFG, German Research Foundation) Germany’s Excellence Strategy EXC 2155 “RESIST” (Project number 39087428), SFB 900/3 (Project B1, 158989968) and FOR2830 (Project FO334/7-1, 398367752). Research in the Messerle lab is funded by DFG Germany’s Excellence Strategy EXC 2155 “RESIST” (Project number 39087428), SFB 900/3 (Project B1, 158989968) and FOR2830 (ME 1102/4-2; 398367752).

## Author contributions

X.L., R.G.J., Y.L., B.B., and R.F. designed the study; X.L., R.G.J., Y.L., S.H., L.O., C.R., A.S., S.W., A.J., and B.B. performed experiments; K.W. and M.M. provided MCMV virus strains; X.L., R.G.J, Y.L., and B.B. analyzed the data; X.L., R.G.J., and B.B. wrote the first draft; B.B. and R.F. supervised the study. All authors have read and agreed to the published version of the manuscript.

## Conflict of interest disclosure

The authors declare no conflict of interest.

## Notes

### Competing Interest Statement

The authors have declared no competing interest.

