## Supplemental Tables and Figures for "Protective function of *ex vivo* expanded CD8 T cells in a mouse model of adoptive therapy for cytomegalovirus infection depends on integrin beta 1 but not CXCR3, CTLA4, or PD-1 expression"

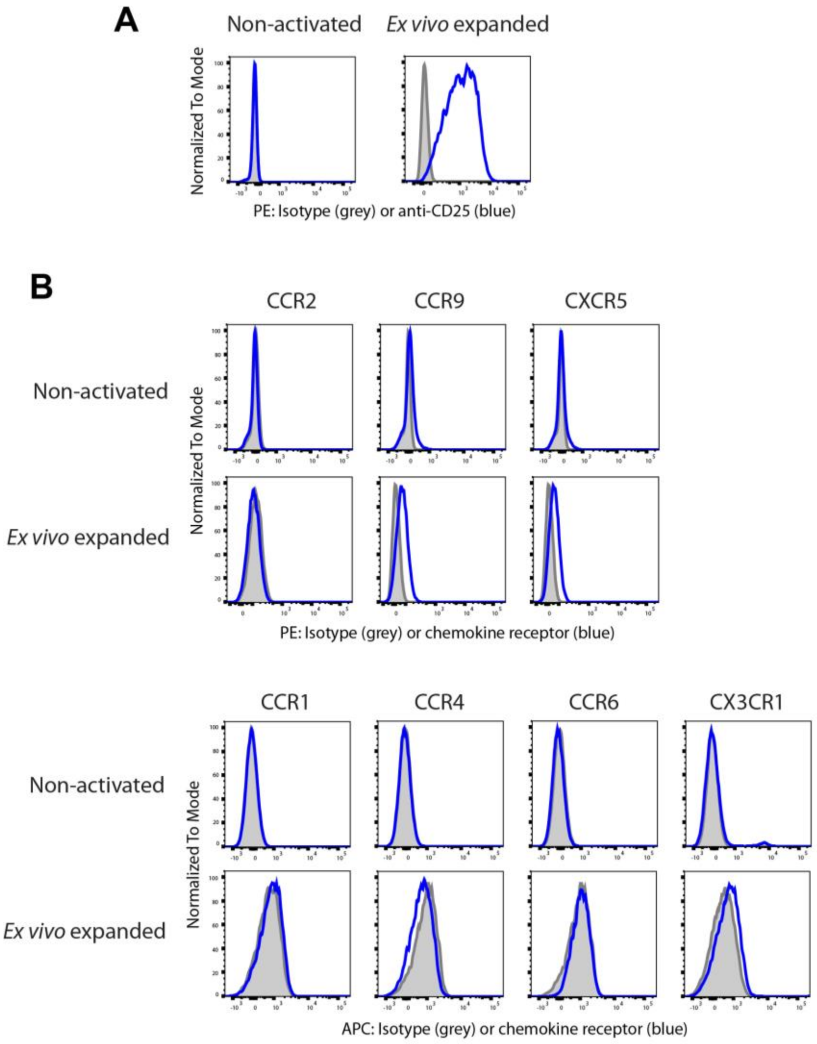


**Supplementary Figure 1. *Ex vivo* expansion alters expression CD25 but not 7 chemokine receptors on OT-I CD8 T cells.** Expression of (**A**) CD25 and (**B**) indicated chemokine receptors on CD8 OT-I T cells before (non-activated) and after 7 days of *ex vivo* expansion: blue lines, antibody staining; gray shaded areas, isotype control. Representative histograms from 2-3 experiments.

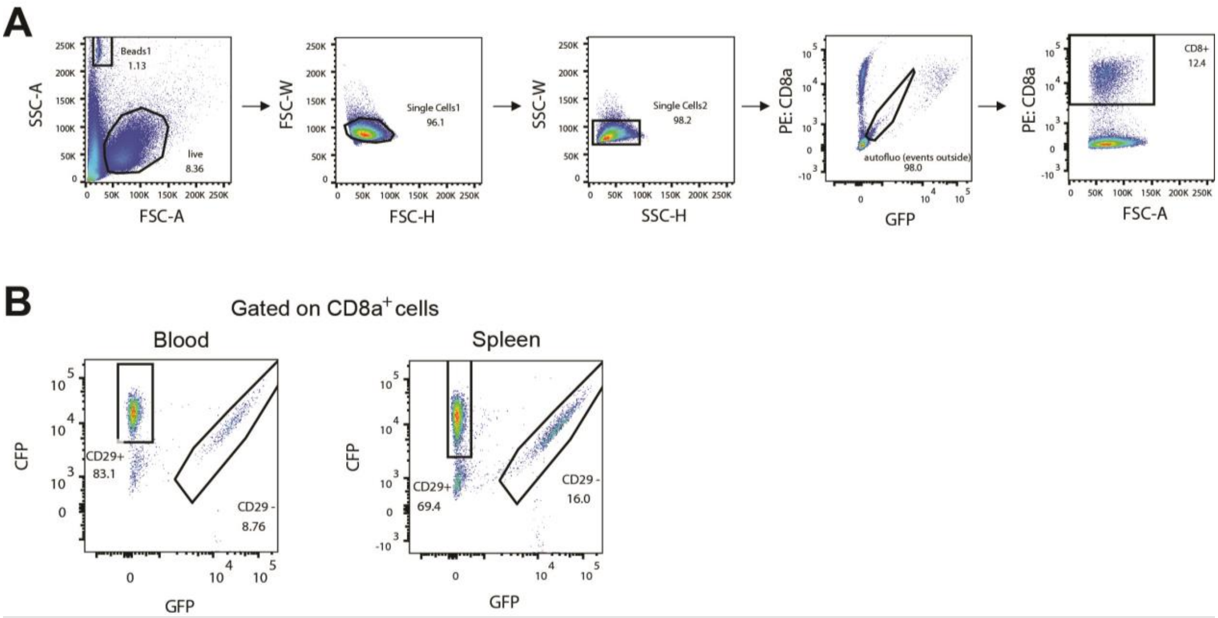


**Supplementary Figure 2. Flow cytometry strategy used to monitor integrin-β1 (CD29)-mediated expansion of antigen-specific CD8 T cells during anti-viral immunotherapy.** (**A**) Gating strategy for adoptively transferred *ex vivo* expanded OT-I cells stained with indicated antibodies (Supplementary Table S3). Pseudocolor plots show representative data from one mouse analyzed at 11 days post infection (dpi) with MCMV-3D and 10 days post OT-I T cell transfer. (**B**) Pseudocolor plots showing ratios of *Itgb1*$^{+/+}$ CFP$^{+}$ and *Itgb1*$^{-/-}$ GFP$^{+}$ OT-I cells at 11 dpi in blood and spleen of a mouse that received $10^4$ of each cell population one day before MCMV-3D intraperitoneal infection. Representative data from 6 analyzed mice in 2 independent experiments are shown.

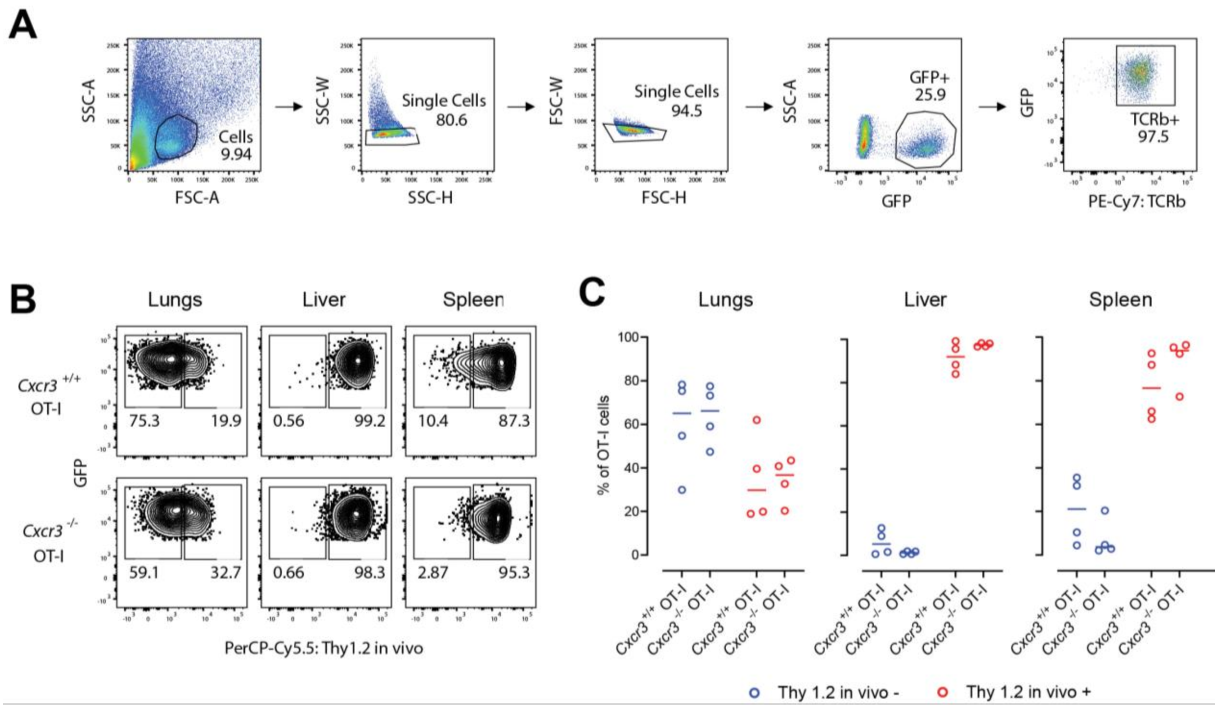


**Supplementary Figure 3. Gating strategy and *in vivo* labeling of OT-I T cells for investigation of CXCR3 role in homing of adoptively transferred *ex vivo* expanded VSTs.** (**A**) Gating strategy for adoptively transferred ex vivo expanded OT-I cells stained with indicated antibodies (Supplementary Table S3). Pseudocolor plots show representative data from one mouse analyzed at 8 days post infection (dpi) with MCMV-3D and 7 days post OT-I T cell transfer. (**B**) Representative contour plots showing staining of $GFP^+$ OT-I cells isolated from indicated organs at 8 dpi with anti-Thy 1.2 antibody that had been intravenously (i.v.) injected 5 min before the mice were sacrificed. Mice received $7.5x10^4$ of either *Cxcr3*$^{+/+}$ $GFP^+$ or *Cxcr3*$^{-/-}$ $GFP^+$ OT-I cells one day before MCMV-3D intraperitoneal infection (N = 4 per group pooled from 4 experiments). (**C**) Distribution of adoptively transferred OT-I T cells in indicated organs according to the *in vivo* staining with anti-Thy 1.2 antibody. Pooled data from 4 experiments, shown as individual mice (dots) and group mean (line).

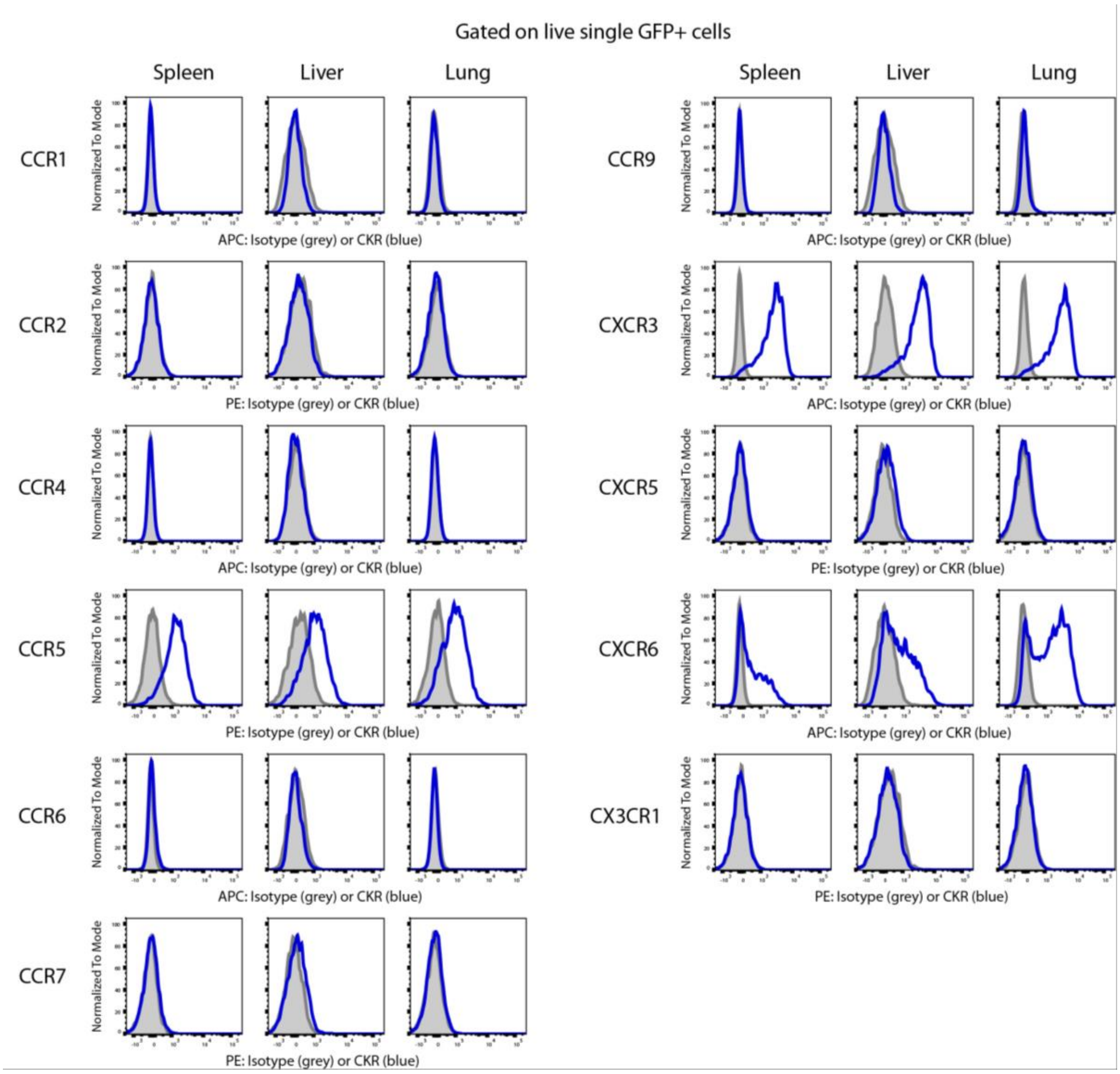


**Supplementary Figure 4. Chemokine receptor (CKR) expression on adoptively transferred OT-I cells isolated from organs of MCMV-3D infected Rag2$^{-/-}$ mice at 8 dpi and 7 days post T cell transfer.** Blue lines, antibody staining; gray shaded areas, isotype control. Representative data of 4 mice from 3 experiments.

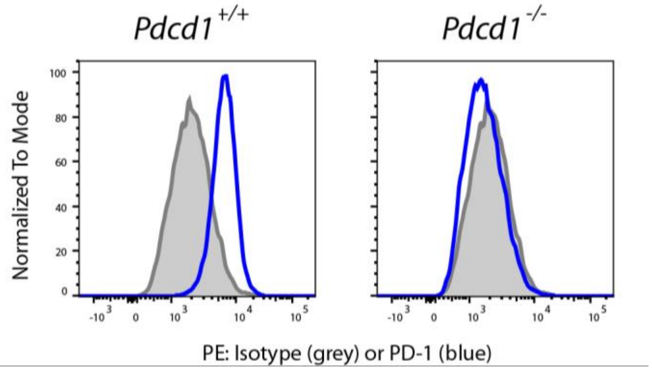


**Supplementary Figure 5. Successful deletion of *Pdcd1* in ex vivo expanded OT-I T cells using CRISPR/Cas9 RNPs.** Representative histograms showing PD-1 surface expression after CRISPR/Cas9-mediated deletion using an negative control gRNA or three different gRNA targeting the *Pdcd1* (N = 5 experiments).

**Supplementary Table 1. List of crRNAs (all from Integrated DNA Technologies Inc; IDT).**

| Gene target | crRNA sequence (5‘->3‘) | crRNA designation |
|---|---|---|
| *Itgb1* | GUGCUUAGUCUUACUGACAG | 02 |
| | AUUACUUCAGACUUCCGCAU | 04 |
| | AGUGACAUAGAGAAUCCCAG | 07 |
| *Cxcr3* | UCUGAACUUCACUCCCACAA | 05 |
| | UGACUCCCCGCCCUGCCCAC | 06 |
| | GCUGUUCUGCUGGUCUCCAG | 07 |
| *Pdcd1* | GGUAUCAUGAGUGCCCUAGU | AC |
| | AGAUCAUACAGCUGCCCAAC | AD |
| | GCAAAAAUCGAGGAGAGCCC | 4 |
| *Ctla4* | AUGGAAAGCUGGCGACACCA | AB |
| | UGGCUUGUCUUGGACUCCGG | AC |

**Supplementary Table 2. List of primers used (all from Sigma)**

| Target gene | primer type | primer sequence (5' -> 3') |
|---|---|---|
| *Pdcd1* | forward primer 1 | CCCCTCTCCCTGTGGAAATC |
| *Pdcd1* | reverse primer 1 | CCACAACACAGGGTAGGCAT |
| *Pdcd1* | forward primer 2 | AACTGCTTACGATATTCTGCCC |
| *Pdcd1* | reverse primer 2 | GCAAGCCCTTCCATCCCTTAAA |
| *Pdcd1* | forward primer 3 | TGAACTGGAACCGCCTGAGT |
| *Pdcd1* | reverse primer 3 | AGAAGCAGGGTATGATGAGCC |
| *Ctal4* | forward primer 1 | TGGAGAGTGACTGACTACAGC |
| *Ctal4* | reverse primer 1 | CCTAGCTATGCATCACAGACC |
| *Ctal4* | forward primer 2 | CGACGTAACAGCTAAACCCA |
| *Ctal4* | reverse primer 2 | GCTCCTTCGCTACTGCTAGAC |

Supplementary Table 1. List of antibodies and other reagents used for cell line characterization and knockout analysis. NA – not applicable

| **Antibody** | **Clone** | **Source** | **Identifier** |
|---|---|---|---|
| *Antibodies used in flow cytometry* | | | |
| FITC hamster anti-mouse CD3ε | 145-2C11 | BioLegend | Cat# 100306, RRID:AB_312671 |
| Brilliant Violet 605™ hamster anti-mouse CD3ε | 145-2C11 | BD Biosciences | Cat# 563004, RRID:AB_2737945 |
| Brilliant Violet 711™ hamster anti-mouse CD3ε | 145-2C11 | BioLegend | Cat# 100349, RRID:AB_2565841 |
| PerCP/Cyanine5.5 rat anti-mouse CD8α | 53-6.7 | BioLegend | Cat# 100734, RRID:AB_2075238 |
| PE rat anti-mouse CD8α | 53-6.7 | BioLegend | Cat# 100708, RRID:AB_312747 |
| Brilliant Violet 510™ rat anti-mouse/human CD44 | IM7 | BioLegend | Cat# 103044, RRID:AB_2561391 |
| Brilliant Violet 785™ rat anti-mouse CD62L | MEL-14 | BioLegend | Cat# 104440, RRID:AB_2629685 |
| PE-Cy7 hamster anti-mouse TCR beta chain | H57-597 | BioLegend | Cat# 109222, RRID:AB_893625 |
| PerCP-Cy5.5 rat anti-mouse CD90.2 (Thy 1.2) | 53-2.1 | BioLegend | Cat# 140322, RRID:AB_2562696 |
| Biotin rat anti-mouse CD18 (Integrin β2) antibody | M18/2 | Thermo Fisher Scientific | Cat# 13-0181-82, RRID:AB_466381 |
| Biotin rat anti-mouse CD29 (Integrin β1) antibody | Ha2/5 | BD Biosciences | Cat# 555004, RRID:AB_395638 |
| Biotin rat anti-mouse Integrin β7 Antibody | FIB504 | Thermo Fisher Scientific | Cat# 13-5867-82, RRID:AB_1518756 |
| Biotin rat anti-mouse CD51 (Integrin αV) Antibody | RMV-7 | Thermo Fisher Scientific | Cat# 13-0512-81, RRID:AB_466476 |
| Biotin rat IgG2a kappa isotype control antibody | 1H4 | Homemade | N/A |
| Biotin rat IgG1 kappa isotype control antibody | CAD9 | Homemade | N/A |
| Biotin hamster IgG isotype control antibody | HTK888 | Biolegend | Cat# 400903, RRID:AB_3094652 |
| Streptavidin APC | N/A | Biolegend | Cat# 405207 |
| Streptavidin Cy5 * | N/A | eBioscience | Cat# 19-4317-82 |
| APC hamster anti-mouse CD152 (CTLA4) antibody | UC10-4B9 | Thermo Fisher Scientific | Cat# 17-1522-82, RRID:AB_2016700 |
| PE rat anti-mouse CD279 (PD-1) antibody | RMP1-30 | BioLegend | Cat# 109104, RRID:AB_313420 |
| APC hamster anti-mouse CD183 (CXCR3) antibody | CXCR3-173 | Thermo Fisher Scientific | Cat# 17-1831-82, RRID:AB_1210791 |
| PE rat anti-mouse CD185 (CXCR5) antibody | SPRCL5 | Thermo Fisher Scientific | Cat# 12-7185-82, RRID:AB_11217882 |
| APC rat anti-mouse CD186 (CXCR6) antibody | SA051D1 | BioLegend | Cat# 151106, RRID:AB_2572143 |
| APC rat anti-mouse CD191 (CCR1) antibody | S15040E | BioLegend | Cat# 152504, RRID:AB_2629811 |

| | | | |
|---|---|---|---|
| PE rat anti-mouse CD192 (CCR2) antibody | SA203G11 | BioLegend | Cat# 150610, RRID:AB_2616982 |
| APC hamster anti-mouse CD194 (CCR4) antibody | 2G12 | BioLegend | Cat# 131212, RRID:AB_2074507 |
| APC hamster anti-mouse CD195 (CCR5) antibody | HM-CCR5 | BioLegend | Cat# 107012, RRID:AB_2074528 |
| PE hamster anti-mouse CD195 (CCR5) antibody | HM-CCR5 | Thermo Fisher Scientific | Cat# 12-1951-82, RRID:AB_657684 |
| APC hamster anti-mouse CD196 (CCR6) antibody | 29-2L17 | BioLegend | Cat# 129814, RRID:AB_1877148 |
| APC rat anti-mouse CD197 (CCR7) antibody | 4B12 | Thermo Fisher Scientific | Cat# 17-1971-82, RRID:AB_469444 |
| PE rat anti-mouse CD197 (CCR7) antibody | 4B12 | Thermo Fisher Scientific | Cat# 12-1971-82, RRID:AB_465905 |
| PE rat anti-mouse CD199 (CCR9) antibody | 9B1 | BioLegend | Cat# 129708, RRID:AB_1227485 |
| APC mouse anti-mouse CX3CR1 antibody | SA011F11 | BioLegend | Cat# 149008, RRID:AB_2564492 |
| APC armenian hamster IgG isotype control antibody | HTK888 | BioLegend | Cat# 400912, RRID:AB_2905474 |
| APC Rat IgG2a kappa isotype control antibody | eBR2a | Thermo Fisher Scientific | Cat# 17-4321-81, RRID:AB_470181 |
| APC rat IgG2b kappa isotype control antibody | RTK4530 | BioLegend | Cat# 400612, RRID:AB_326556 |
| PE rat IgG2a kappa isotype control antibody | eBR2a | Thermo Fisher Scientific | Cat# 12-4321-42, RRID:AB_1518773 |
| PE rat IgG2b kappa isotype control antibody | eB149/10H5 | Thermo Fisher Scientific | Cat# 12-4031-82, RRID:AB_470042 |
| *Antibodies used for immunofluorescent microscopy* | | | |
| APC rat anti-mouse CD45 antibody | 30-F11 | BD Biosciences | Cat# 559864, RRID:AB_398672 |

* this product is discontinued
